# Characterization and Validation of a Middle-Down Hydrophobic Interaction Chromatography Method to Monitor Methionine Oxidation in IgG1

**DOI:** 10.1101/2023.02.01.525715

**Authors:** Somar Khalil, Nisha Patel, Francoise Bevillard-Kumar, Cyrille Chéry, William Burkitt, John O’Hara, Annick Gervais

## Abstract

Post-translational modifications (PTMs) of therapeutic monoclonal antibodies (mAbs) can impact the efficacy of a drug. Methionine oxidation can alter the overall hydrophobicity of an antibody, thereby inducing conformational changes and affecting its biological activity. To ensure high quality, safety, and efficacy of mAbs, routine monitoring of PTMs such as methionine (Met) oxidation is essential. Met oxidation in the fragment crystallizable (Fc) region of immunoglobulin-G1 (IgG1) is a critical quality attribute because it impacts not only the interaction with the neonatal Fc receptor and protein A but also the half-life of mAbs in serum circulation. Although bottom-up mass spectrometry provides high site specificity, it may have limited application in quality control workflows, and its complicated sample preparation could result in procedure-induced oxidation. In this study, we describe the development and characterization of a rapid and robust middle-down hydrophobic interaction chromatography method for monitoring Met oxidation in the Fc region of IgG1. Additionally, we assessed a comprehensive method validation package and demonstrated the specificity, linearity, precision, and accuracy of the new method within a range of 3.8–37.7%. The relative quantitative data provided by this method may be used in a regulated workflow to support process and formulation development as well as in the later stages of drug development and batch release and stability studies.

## Introduction

Monoclonal antibodies (mAbs) are widely recognized therapeutic agents owing to their high biological activity and favorable physicochemical characteristics^1^. The expanding number of licensed antibody medications and promising clinical candidates demonstrate that developing mAbs as therapeutics is a primary focus for the scientific community^2-4^. Because of the complex structures of mAbs, the impact of chemical modifications on their physical stability and aggregation remains an area in need of further investigation^2,5,6^. Moreover, maintaining structural and chemical integrity during drug development is key to ensuring efficacy and safety^7^.

Oxidation is a commonly observed chemical degradation pathway in mAbs^8,9^. It can alter the hydrophobicity of a protein surface, affecting its conformation and thus its biological activity^10-13^. Moreover, the oxidation of mAbs impairs their stability^14^, and degradation pathways such as fragmentation and aggregation have been reported to become more susceptible as a result of oxidation^15,16^. Methionine (Met), tryptophan (Trp), histidine (His), lysine (Lys), and cysteine (Cys) are major amino acid residues in proteins that are susceptible to oxidation^17-19^; the oxidation reaction of Met is considerably faster than that of the other residues^17,20,21^. Immunoglobulin-G1 (IgG1) molecules have two conserved Met residues: Met252 in the CH2 domain and Met428 in the CH3 domain (EU numbering). Both are highly vulnerable to oxidation^22^, and the impact of oxidation on both residues in fragment crystallizable (Fc)– neonatal Fc receptor (FcRn) binding has been extensively studied in vitro and in vivo^8,19,23-25^. It has been identified that Met oxidation in the Fc weakens the binding affinity of IgGs to protein A and FcRn^23,26,27^. The oxidation of each Met252 residue on both heavy chains has been shown to impact the half-life of mAbs in serum circulation^26^.

Ion exchange chromatography (IEX) is an established technique routinely used to monitor the charge profile of mAbs^28,29^. Since it has been reported that oxidized mAbs are basic in nature, it is an attractive option for monitoring Met oxidation. However, mAbs are inherently complex owing to other basic post-translational modifications (PTMs) such as Lys clipping, isomerization, and succinimide formation. Therefore, specific monitoring of Met oxidation by IEX is challenging. Monitoring oxidation via FcRn binding can also be achieved by studying ligand interactions, for example, via surface plasmon resonance^8,24,30^ or affinity chromatography^27^. The main drawbacks of routinely using ligand chromatography are the cost of buying commercial FcRn columns or ensuring batch-to-batch consistency of home-built affinity columns.

Bottom-up analysis using liquid chromatography coupled with tandem mass spectrometry (LC-MS/MS) is the most specific and accurate method for the monitoring of most PTMs^31,32^. However, this approach requires lengthy sample preparation, expensive instrumentation, and complicated analysis. The bottom-up analysis involves intricate and relatively low throughput sample preparation in which oxidation can be further induced and may restrict its application in quality control (QC) workflows^33-36^. Alternatively, the middle-down approach easily generates IgG1 sub-domain fragments by special proteases under native conditions, avoiding the need for time-consuming sample preparation and digestion as well as the potential PTM information loss associated with bottom-up analyses^37-43^.

A substantial part of the nonpolar residues in mAbs are exposed to water interfaces, which is contrary to the preferential positioning of hydrophobic amino acids in the interior of the biomolecule^44,45^. Such nonpolar amino acids generate patches of unique hydrophobic character on the protein surface, which could account for the protein’s bio-specific conformation and its propensity to form complexes or aggregates^46,47^. Hydrophobic interaction chromatography (HIC) is a milder form of reversed-phase liquid chromatography (RPLC), exploiting the hydrophobic properties of the biomolecules to separate them^48,49^. In HIC, species of interest containing hydrophobic and hydrophilic regions are applied to a column in a high-salinity buffer. The salt in the buffer helps to prevent sample solutes from solvating, and the hydrophobic regions are adsorbed by the stationary phase. The samples are then eluted from the column using a decreasing salt gradient in the order of increasing hydrophobicity.

This study aimed to develop a sensitive and robust middle-down HIC method for the rapid separation and relative quantification of oxidized Met in the Fc region of IgG1-type mAbs, making it ideal for the biopharmaceutical industry. To aid in the development of the method, a combination of mass spectrometry (MS) techniques was employed to characterize proteoforms found across the entire HIC profile. Furthermore, performing size-exclusion chromatography (SEC)-MS analysis allowed us to investigate the viability of HIC for structural variant separation. After the peaks containing oxidized Fc were identified, the method was subsequently developed and validated in accordance with the International Council on Harmonization (ICH Q2) guideline for impurity testing (specificity, linearity, precision, accuracy, quantitation limit (QL), and robustness).

## Materials and Methods

### Materials

A humanized, full-length IgG1-type mAb with a molecular weight of approximately 150 kDa was used. Chemically oxidized mAb was generated by incubation with H_2_O_2_. Incubation was carried out with 0.1% (w/w) H_2_O_2_ for 14 d at 5 °C or with 0.01% (w/w) H_2_O_2_ for 72 h at 5 °C for characterization by MS or for HIC analysis, respectively. The reaction was quenched by buffer exchange into formulation buffer. Temperature-stressed mAb was generated by incubation for 180 d at 25 °C. Protein-Pak HIC (4.6 × 100 mm, 2.5 μm), BEH C18 (2.1 × 150 mm, 1.7 μm) and BEH SEC (4.6 × 150 mm, 1.7 μm) columns were purchased from Waters corporation. MAbPac HIC (4.6 × 100 mm, 5 μm), dithiothreitol (DTT) and iodoacetamide (IAM) were purchased from Thermo Scientific. AdvanceBio HIC (4.6 × 100 mm, 3.5 μm) column was purchased from Agilent Technologies. FabRICATOR and FabALACTICA enzymes were purchased from Genovis. Sequencing grade trypsin and Ides protease were purchased from Promega. Papain enzyme, guanidine hydrochloride, ammonium acetate, ammonium sulfate, sodium sulfate, sodium chloride and H_2_O_2_ were purchased from Sigma-Aldrich. Acetonitrile (ACN), isopropyl alcohol (IPA), trifluoroacetic acid (TFA), and formic acid (FA) were purchased from Biosolve. Phosphate buffer solution (PBS) was purchased from Invitrogen.

### Method Development

#### Stationary phase

In contrast to RPLC, weakly hydrophobic ligands such as short-chain alkyl and phenyl groups are used on the packing material in HIC^50^, resulting in milder protein–ligand hydrophobic interactions. For bio-separations by HIC, both the chemistry and density of the ligands are critical because they can alter the HIC selectivity and adsorption capacity^51,52^. In this study, three stationary phases with distinctive characteristics were assessed. As presented in Fig. S3, the porous columns with polyalkylimide ligands (MabPac) or with proprietary ligands^53^ (AdvanceBio) exhibited superior resolution between peaks when compared with the non-porous column that contains butyl ligands (Protein-Pak). Showing less peak tailing, the AdvanceBio column was selected for this study.

#### Mobile phase

Various mobile phase elements, such as salt type and concentration, pH, and the presence of organic modifiers, can potentially impact bio-separation by HIC^54-58^. Following the Hofmeister series^59^, a neutral salt (sodium chloride) and anti-chaotropic salts (ammonium and sodium sulfate) were investigated at different concentrations. Ammonium sulfate at 1 M concentration was selected as it provided the best resolution. Changes in mobile phase pH may alter the selectivity by impacting the degree of hydrophobic interactions according to the pI and number of charged amino acid residues in the mAb^57,60^. pH values near physiological conditions were evaluated and a pH of 7.0 was selected for this study. Adding a small amount of organic modifier to the mobile phase may also alter the selectivity by disrupting the conformational structure of the mAb and exposing more hydrophobic areas that can interact with the stationary phase^58^. We experimented with both ACN and IPA, and while adding 5% ACN marginally improved the resolution, we chose not to add organic modifiers to the mobile phase in order to maintain non-denaturing conditions.

#### Protease enzyme

The mAb hinge region can be targeted by a middle-down approach with special proteases^61-63^. In this study, we evaluated the specific below-the-hinge cleaving enzyme IdeS, which yields F(ab’)2 and Fc fragments^38^. Papain, a Cys protease that cleaves the mAb above the hinge to yield two monovalent Fab fragments and one intact Fc/2 fragment^64,65^, was also investigated. To induce the catalytic activity of papain, reducing conditions are required to reduce the thiol group of the Cys25 residue in the enzyme^66,67^. Finally, this study evaluated IgdE, a Cys protease derived from *S. agalactiae* under the commercial name FabALACTICA, which cleaves the mAb above the hinge in native conditions^64^ (Fig. S4). Interestingly, both IdeS and IgdE enzymes provide high specificity for the hinge region of the mAb and prevent the under- and over-digestion that is common with papain^68^. Ultimately, we selected IdeS digestion for this study because a primary objective was to monitor Fc Met oxidation. Therefore, generating Fc fragments was more favorable.

### Middle-Down HIC Method

All samples were diluted with PBS (pH 7.0) to a final concentration of 5.0 mg/mL. The solutions were subsequently digested with an IdeS or a FabRICATOR enzyme at 37 °C for 30 min. The ratio of protein to enzyme was 2:1 (w/unit). Digestion was then quenched by addition of mobile phase A (1 M ammonium sulfate and 50 mM sodium phosphate, pH 7.0). An ACQUITY UPLC H-Class Bio system (Waters) equipped with a TUV detector and AdvanceBio column was employed. The mobile phase A was used for the interaction, and the mobile phase B (50 mM sodium phosphate, pH 7.0) was used for elution. The UPLC gradient was 0–100% B for 14 min at a flow rate of 0.5 mL/min, with a total run time of 22 min. The column oven temperature was set to 30 °C, and the autosampler temperature was set to 5 °C. The UV signal was monitored at 220 nm. Data acquisition and processing were carried out using Empower3 (Waters).

### Collection of HIC Fractions

Fractions were collected using a shortened HIC method on a UPLC system with no UV detection. Fractions were obtained from more than 40 injections, and then pooled, concentrated, and dialyzed into the native mAb formulation buffer.

### Bottom-Up LC-MS/MS Analysis of mAb and HIC Fractions

mAb samples and fractions were denatured in 8 M guanidine hydrochloride and reduced with 16 mM DTT at 37 °C for 30 min. Alkylation with 44 mM IAM was carried out in the dark at ambient temperature for 30 min. The alkylated samples were then digested with 1:14 (w/w) trypsin at 37 °C for 3 h. The reaction was acidified with 0.065% TFA. Peptides were separated at 40 °C on a BEH C18 column using a Thermo Vanquish high-performance liquid chromatography (HPLC) instrument. Samples were injected at 0.2 mL/min flow rate. Mobile phase A consisted of 0.1% FA in water and mobile phase B consisted of 0.1% FA in ACN. A gradient of 1 to 50% B in 32 min was applied, followed by a 5 min column wash at 99% B. The HPLC instrument was coupled with a Q-Exactive Biopharma MS (Thermo). Tandem MS analysis was performed, and the resulting data were processed and searched against the sequence of the mAb using Biopharmafinder v3 (Thermo). A fixed modification of carbamidomethylated cysteine was specified, and standard modifications typically observed on antibodies were set as variable modifications. The processed data were exported to Excel, and the PTM levels were calculated as follows:

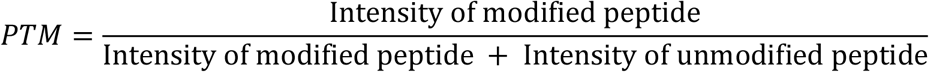

This procedure was performed for each class of modification; for example, Met sulfoxide and Met sulfone levels were individually calculated. Then, both modifications were summed up to provide the overall level of Met oxidation products at a single residue.

### SEC-MS Analysis of HIC Fractions

SEC-MS was performed using a Vanquish HPLC instrument equipped with a BEH SEC column. Isocratic elution using 100 mM ammonium acetate (pH 6.8) was applied at 0.3 mL/min for 20 min. Intact masses were recorded using a Q-Exactive Biopharma MS in high mass range mode (1500–8000 m/z) at a resolution of 35,000 full width half maximum. In-source dissociation was 80 eV. Raw data were deconvoluted to molecular masses using Byos v3.7 (Protein Metrics).

## Results

### MS-Based Characterization

#### Bottom-up LC-MS/MS analysis of native and H_2_O_2_-stressed mAbs

PTM levels were measured in native and hydrogen peroxide (H_2_O_2_)-stressed mAb samples by bottom-up LC-MS/MS analysis (Table 1). Met252 and Met428 are considered to be the primary oxidation events in the Fc region because of their comparatively high solvent accessibility, and their oxidation is associated with impaired IgG binding affinity to FcRn^26^. Native mAb levels of Met252 and Met428 oxidation were 5.7% and 3.1%, respectively. These levels were above 99% in the H_2_O_2_-stressed mAb, and Met sulfoxide was the predominant oxidation product (addition of a single oxygen atom). This indicated that Met252 and Met428 on both heavy chains were almost completely oxidized in H_2_O_2_-stressed mAb. Conversely, the three Met residues in the F(ab’)2 region were responsible for a total level of oxidation of 5.7% in the native mAb and 17.4% in the H_2_O_2_-stressed mAb. H_2_O_2_-induced oxidation is expected to largely affect the Met oxidation levels with minimal impact on other residues such as His and Trp^20,69^. Indeed, His oxidation did not significantly increase after H_2_O_2_ treatment, whereas Trp oxidation slightly increased by ∼ 2.5-fold (Table 1).

**Table 1.** Summary of % PTMs in native and H_2_O_2_-stressed mAbs analyzed by bottom-up LC-MS/MS.

| **PTM (%)** | **Native mAb** | **H<sub>2</sub>O<sub>2</sub>-stressed mAb** |
| --- | --- | --- |
| Met252 oxidation | 5.7 | 99.2 |
| Met428 oxidation | 3.1 | 99.1 |
| Fc Met oxidation | 16.5 | 100 |
| Fc Trp oxidation | 6.5 | 17.4 |
| Fc His oxidation | 0.6 | 1.4 |
| C-terminal lysine clipping | 97.2 | 96.9 |
| N-terminal pyroglutamic acid | 1.5 | 1.1 |
| F(ab')<sub>2</sub> Met oxidation | 5.7 | 17.4 |
| F(ab')<sub>2</sub> Trp oxidation | 6.2 | 16.8 |
| F(ab')<sub>2</sub> His oxidation | 0.9 | 3.5 |

#### Characterization of HIC fractions

Native and H_2_O_2_-stressed mAb samples were subjected to immunoglobulin-degrading enzyme of *Streptococcus pyogenes* (IdeS) digestion and fractionated across a shortened HIC method into eight fractions (Fig. 1). The fractions were subsequently checked for purity by HIC and characterized by bottom-up LC-MS/MS and SEC-MS analysis. The eight fractions were representative of a single predominant peak as well as adjacent peaks. Met, His and Trp oxidation were reported along with C-terminal Lys clipping and N-terminal pyroglutamic acid.

**Figure. 1.**
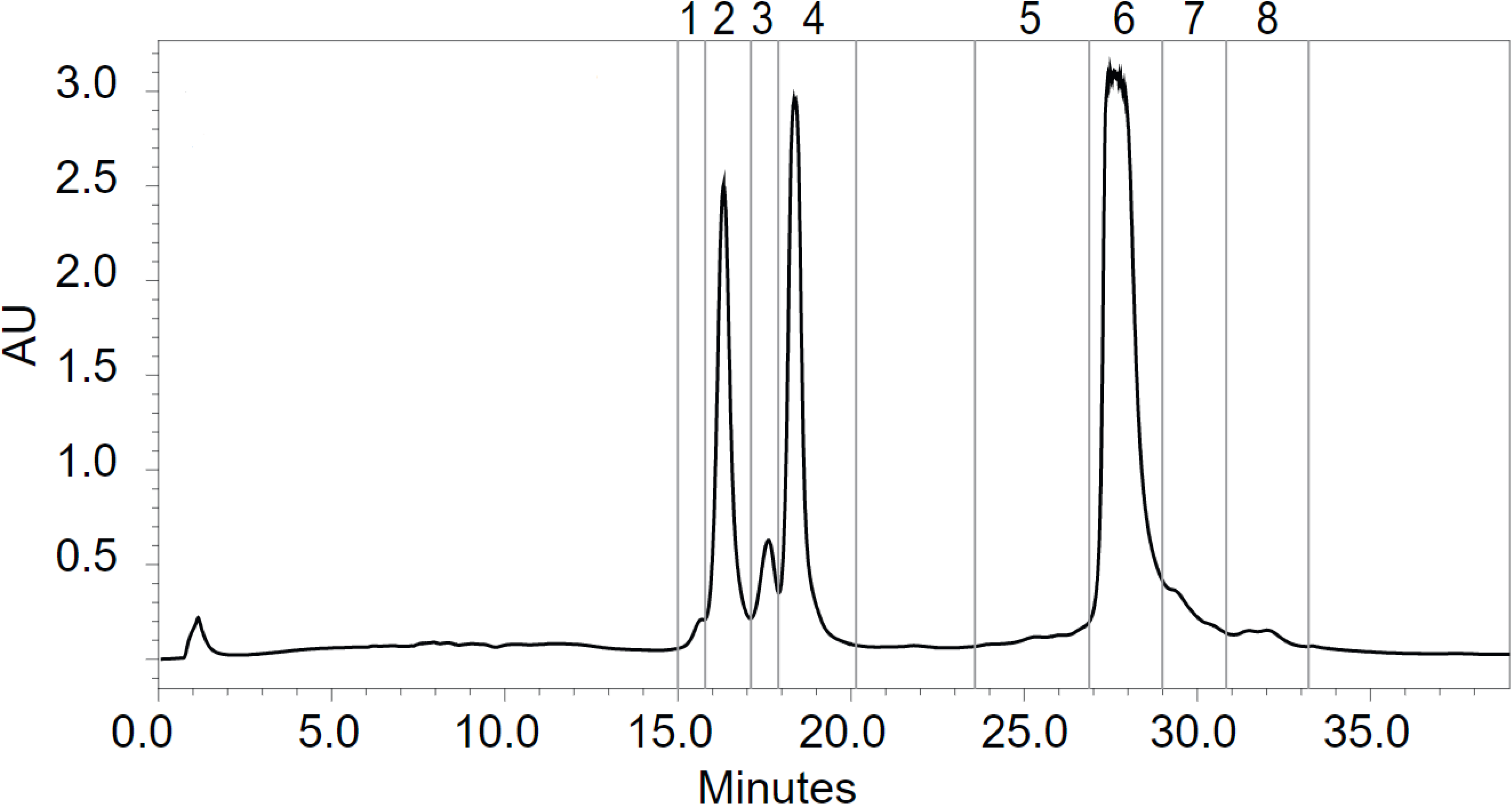
Ides-digested mAb fractions obtained from a shortened HIC method. Eight fractions were created using a shortened HIC method after IdeS digestion of the native and H_2_O_2_-stressed mAb samples.

#### Characterization of Fc-containing fractions

The Fc subunit in the early-eluting fractions (1-4) ranged from 98.7-99.8% of the protein composition (Table S1). Met252 and Met428 were both nearly fully oxidized in fractions 1 and 2, corresponding to Fc with quadruple oxidation variant (Met252/Met252 and Met428/Met428). Clipped C-terminal Lys variants attributed to the difference between the two fractions (63.3% in fraction 1 vs. 97.8% in fraction 2) corresponding to variants with one Lys clipped (peak A), and both Lys clipped (peak B) (Fig. 2). Fraction 3 was found to contain a mixture of peaks which was comprised of one main isoform and other isoforms from adjacent peaks (Table S1). The isoforms in fraction 3 were distinguished by SEC-MS (Fig. S1), and the elution order of these isoforms was inferred by using supportive evidence from adjacent fractions. In summary, the double oxidation variant (Met252/Met252, Met428/Met428, or Met252/Met428) with fully clipped Lys was attributed to peak C, and two isoforms were attributed to peak D; singly oxidized variant (Met252 or Met428) with both C-terminal Lys clipped and a non-oxidized variant with one C-terminal Lys clipped. Fraction 4 corresponded to Fc with no oxidation events and fully clipped C-terminal Lys (peak E).

**Figure. 2.**
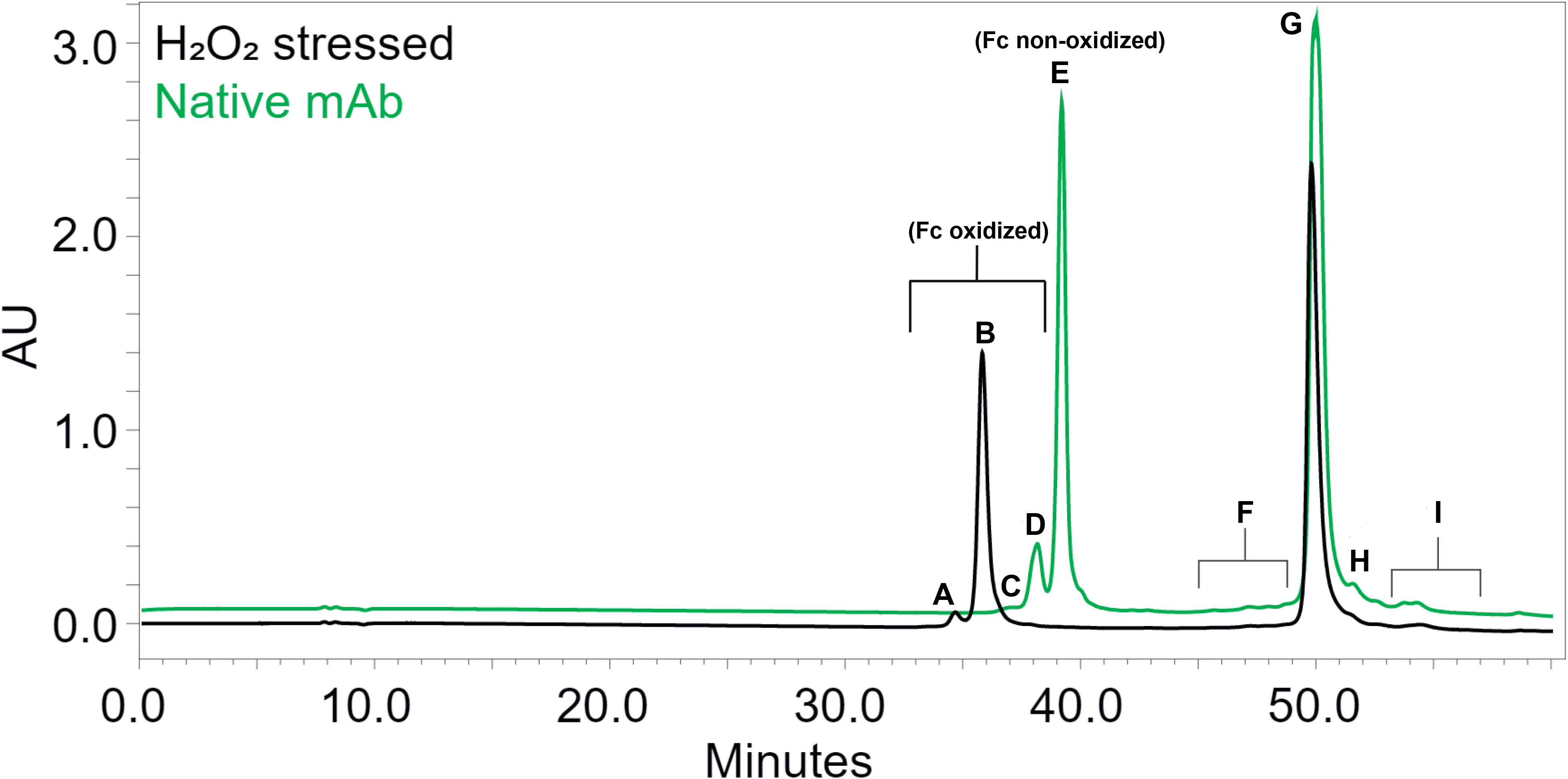
Fc and F(ab’)2 species in HIC chromatograms of Ides-digested native and H_2_O_2_-stressed mAb samples. HIC chromatograms of Ides-digested native (green) and H_2_O_2_-stressed (black) mAb samples. Peaks A, B, C, and D combined represent Fc oxidized. Peak E represents Fc non-oxidized. Peaks F, G, H, and I combined represent F(ab’)2.

#### Characterization of F(ab’)2-containing fractions

The late-eluting fractions (5-8) were predominantly composed of bivalent fragment-antigen binding (F(ab’)2) subunits (Table S2). Owing to their larger size, F(ab’)2 isoforms were not as well resolved as the Fc isoforms. Low-level variants of F(ab’)2 isoforms, Fc, and noncovalent complexes made up fraction 5. The noncovalent complexes included a F(ab’)2 homodimer and a F(ab’)2/Fc heterodimer. F(ab’)2 isoforms constituted a variant with three truncated N-terminal residues and a mixture of PTMs (as indicated by the mass difference of +11 Da from the theoretical mass). Interestingly, the F(ab’)2/Fc heterodimer complex was detected at low levels across the late-eluting fractions (5, 7, and 8), and while the data quality was not sufficient to assign PTMs that may have caused the F(ab’)2/Fc to elute over several fractions, this behavior could be explained by their multiple conformations. As expected, fraction 6 was comprised solely of F(ab’)2 with lower levels of PTMs. Fraction 7 was also predominantly F(ab’)2 with an enrichment of pyroglutamic acid. Fraction 8 contained low levels of the F(ab’)2/Fc heterodimer, while the predominant species was once again F(ab’)2. By employing SEC–MS, we were able to resolve several compact sub-populations of F(ab’)2 conformations. (Fig.S1).

#### Calculation of Fc Met oxidation levels

In the mAb used in our study, Met oxidation is the predominant event in the Fc region compared to Trp and His oxidation. Trp oxidation levels were ∼15-fold lower than Met oxidation levels in the tested mAb which is likely representative of other standard IgG molecules. Therefore, this HIC method is deemed suitable for monitoring the overall level of Fc Met oxidation in IgGs by integrating the early-eluting Fc-containing peaks that were identified for having single, double, or quadruple Met oxidation events. For the remainder of the study, Fc oxidation will be monitored as a single species (sum of peaks A, B, C and D) and termed “Fc oxidized” (Fig. 2). The ratio of Fc oxidized to the sum of all Fc species (Fc oxidized species + peak E) was used to determine the level of Met oxidation.

### Validation of the HIC Method

To support further application of the HIC method as a stability-indicating assay in the QC environment, this method was validated as an impurity test based on the parameters described in the ICH guideline Q2 (ICH-Q2) for Phase III. The specificity of the method was demonstrated by analyzing the formulation buffer and the mAb. At the retention time of the Fc species, no interference from formulation buffer components was observed. Using E-noval v4.1 (Pharmalex), the total error approach was applied to compute the accuracy profile and predict the quality of the future results with a risk of 5%. The linearity, recovery, precision, accuracy, range, and QL were deduced from the accuracy profile. The linearity was evaluated by analyzing six levels (25, 50, 70, 100, 150, and 250% of the target injected load). The experimental levels (relative %Fc oxidized, normalized by level) were plotted against their theoretical levels (mean plot of the triplicate preparations at each level). The assay deemed linear with coefficient of determination (r^2^) = 1.00 with no visible pattern in the residuals plot. Fc oxidized recoveries were 98.8% and 101.3%. The assay demonstrated repeatability (%RSD of 0%) and precision (%RSD between 1% and 3%). The accuracy of the method was assessed by linking the lower and upper bounds of the tolerance interval calculated for %Fc oxidized. The acceptance limits of the β-tolerance intervals were set at ±30% such that each future result has a 95% chance to fall within (β = 0.95) (Fig. S2). The method proved to be linear, accurate, and precise within the range of 3.8–37.7% Fc oxidized, and the QL was 3.8%, corresponding to the lowest level of the tested range for which the linearity and accuracy criteria were satisfied. To demonstrate stability-indicating characteristics and the ability of the assay of detecting changes in %Fc oxidized in the mAb upon stress, native and temperature-stressed mAb samples were analyzed in triplicates and the mean values (n = 3) of the %Fc oxidized obtained in both types of samples were used to calculate the absolute difference as follows:

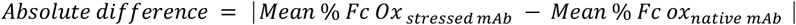

A visual difference between the profiles of native and temperature-stressed mAb samples was observed (Fig. 3), and the calculated absolute difference was greater than the differences observed during intermediate precision (RSD_IP_) at the 100% target injected load.

**Figure. 3.**
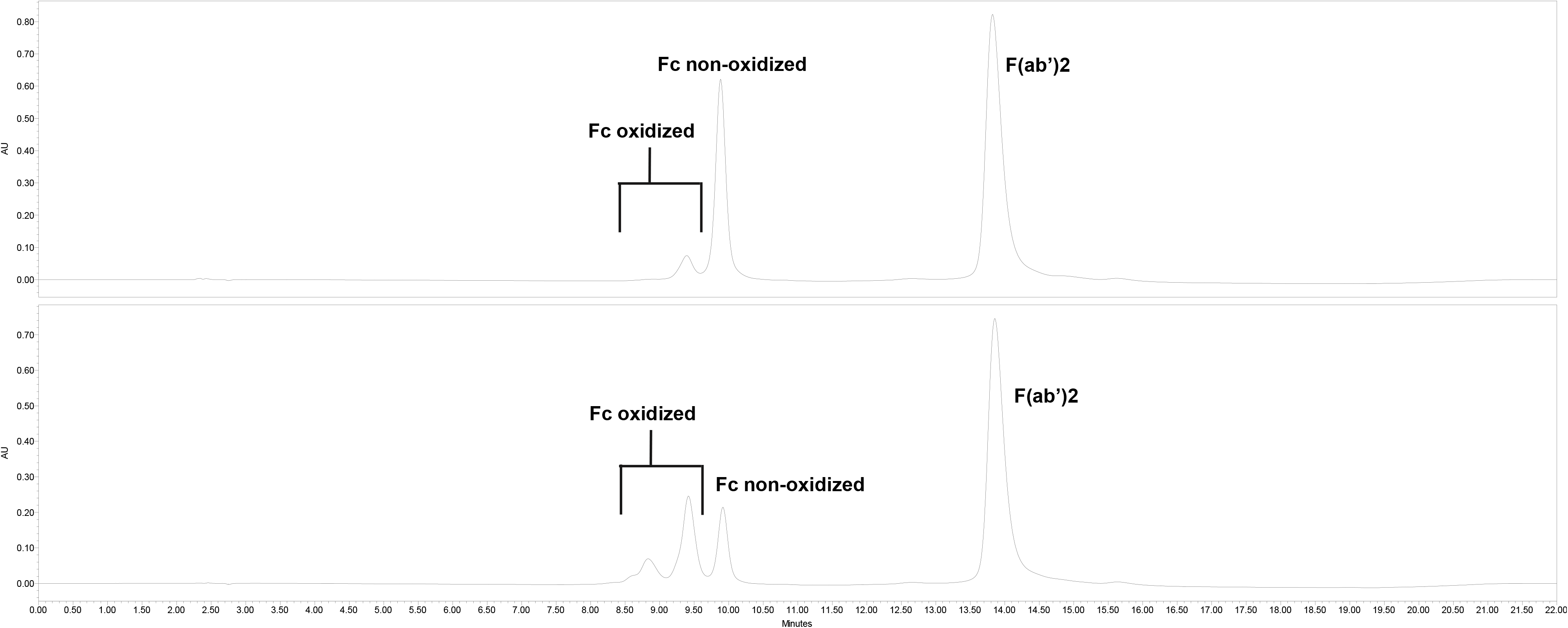
HIC UV chromatograms of Ides-digested native and temperature-stressed mAb samples. A comparison between the HIC chromatograms of Ides-digested native (a) and temperature-stressed (b) mAb samples.

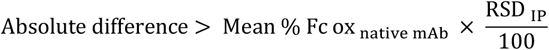

The robustness of some analytical parameters of the HIC assay was assessed following the design of experiment (DoE). Sample preparation parameters (the volume of the digesting enzyme, incubation time, pH of the digestion buffer, and incubation temperature) and chromatographic parameters (the salt concentration of mobile phase A, pH of mobile phase B, and column temperature) were identified as potentially critical. A 20-run split-plot custom design for sample preparation DoE and a 16-run split-plot custom design for chromatographic DoE. In each DoE, four center points for testing all main effects and a two-way interaction between the parameters. For both DoEs, a statistical analysis was performed at a significance level of 5% (α = 0.05), and a step-by-step backward elimination procedure was used to exclude the parameters that were not significant. All sample preparation and chromatographic parameters were deemed robust with no significant effect on the response variable. Incubation time was the only statistically significant parameter (*p*-value = 0.0377). However, this was analytically irrelevant, as the range of values obtained for this parameter was within the accepted range based on the analytical variability which was calculated as follows:

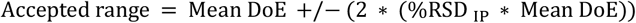

## Discussion

The development of a middle-down HIC approach for the generation of Fc and F(ab’)2 subunits and further resolution of oxidized Fc variants was accomplished by applying a systematic approach based on the principles of HIC and the biophysical characteristics of the mAb used in this study. IdeS was the most appropriate enzyme to generate Fc fragments for measuring Met252 and Met428 oxidation. Salt composition, gradient, and choice of stationary phase were the principal factors contributing to the development of the middle-down HIC method. Our method showed robustness and reliability in monitoring subtle Met252 and Met428 oxidation events in the Fc domain of IgG1s. A minor overestimation of the overall Fc Met oxidation owing to the Lys-clipped variants and Trp oxidation might be observed in mAbs that are subjected to conditions that are known to induce Trp oxidation or change processing of C-terminal Lys, and in-depth characterization by MS would be warranted to determine if the changes are attributed to Met or Trp oxidation or C-terminal Lys clipping. Furthermore, the method demonstrated the capacity to resolve mAb variants with termini modifications and the possibility to study higher-order structure.

Owing to the nature of the enzyme used in our method, which cleaves below the IgG1 hinge region and thus produces larger F(ab’)2 units, it will be challenging to monitor Met oxidation in the fragment-antigen binding (Fab) domain without considerably increasing the gradient time. A targeted approach to monitoring oxidation levels in the Fab domain, however, could be established using enzymes such as FabALACTICA (IgdE), which cleaves at a specific site above the IgG1 hinge region. Along with an intact Fc/2 and a hinge dimer, this enzyme also generates monovalent Fab fragments, which may be better suited for HIC analysis than F(ab’)2 fragments. A monovalent Fab will also reduce the number of structural isoforms associated with the flexible F(ab’)2, facilitating the detection of primary structural isoforms. Furthermore, to extend this method to the analysis of IgG4s, further development would be required owing to the presence of an additional Met residue in the Fc region.

## Conclusions

With the rapid and consistent development of therapeutic antibodies, there is a greater emphasis on understanding the critical quality attributes (CQAs) of mAbs to develop therapeutics with enhanced safety and efficacy. Several studies have indicated that Met oxidation impacts the biological efficacy, safety, and immunogenicity of mAbs, suggesting that it should be monitored as CQA during the discovery, development, and manufacturing processes. In comparison with bottom-up LC-MS/MS, middle-down HIC has fewer time-consuming steps and achieves separation under non-denaturing conditions leading to fewer procedure-induced artifacts that could interfere with accurate oxidation-level assessment in mAbs. Our method can serve as an effective tool for monitoring Met oxidation in the Fc domain of IgG1s. The method was validated to be suitable for routine workflow in QC. It could be applied to support upstream and downstream processes, formulation development, and processes that occur throughout the latter stages of the drug development lifecycle.

## Acknowledgements

We would like to thank Florence Lago, Aurélien Dennel, and Florian Verna for their contributions to the development and validation experiments, as well as Kieran Dawkins for his characterization work.

