## Supplementary material for "Characterization and Validation of a Middle-Down Hydrophobic Interaction Chromatography Method to Monitor Methionine Oxidation in IgG1": Fig. S4

**Supplemental Figure S4:** Middle-down HIC analysis of H_2_O_2_-stressed mAb. (A) after Papain digestion. (B) after IdeS digestion. (C) after Fabalactica digestion.

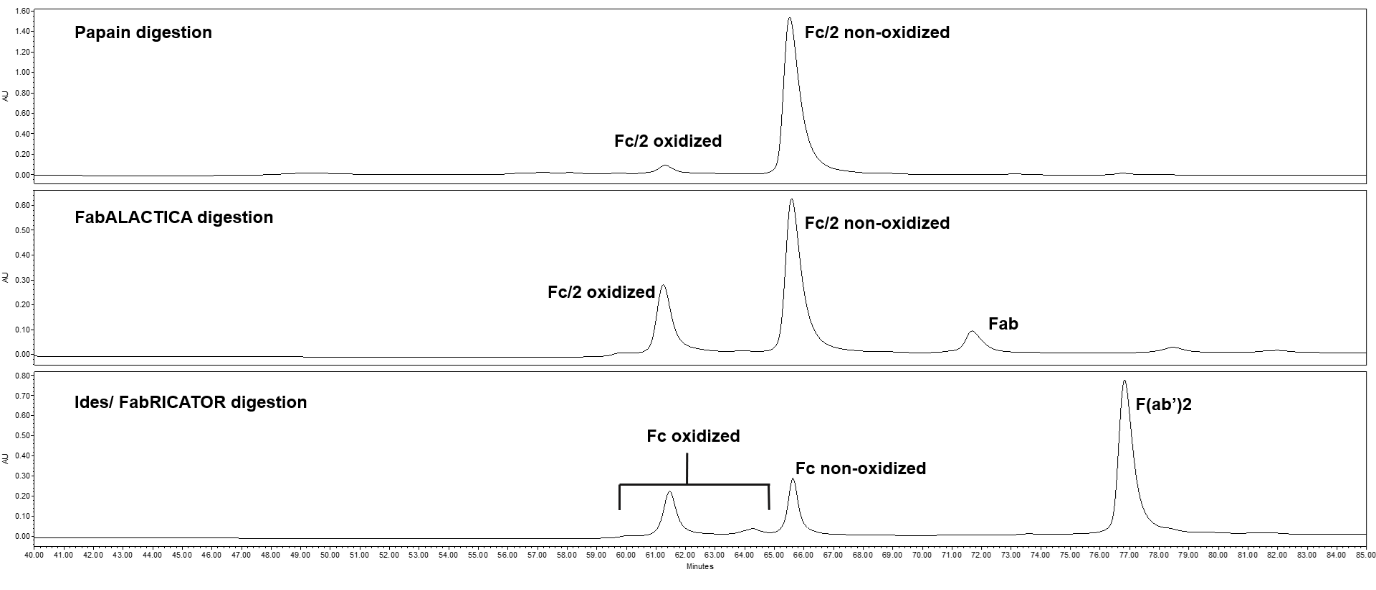
