## Supplementary material for "Characterization and Validation of a Middle-Down Hydrophobic Interaction Chromatography Method to Monitor Methionine Oxidation in IgG1": Table S1

**Supplemental Table S1:** Summary of % PTMs, bottom-up LC-MS/MS and SEC-MS analysis of purified Fc-containing fractions. Bold indicates predominant species.

| Fraction | F(ab’)2:Fc (%) | M252 oxidation (%) | M428 oxidation (%) | Fc Met oxidation (%) | C-terminal Lysine clipping (%) | Fc Trp oxidation (%) | Fc His Oxidation (%) | SEC-MS |
| --- | --- | --- | --- | --- | --- | --- | --- | --- |
| 1 | 1.3: 98.7 | 97.7 | 98.9 | 100 | 63.3 | 16.4 | 1.6 | **Fc, 4 oxidation,**  **-K/unclipped** |
|  |  |  |  |  |  |  |  | Fc, 4 oxidation, -K/-K |
| 2 | 0.2: 99.8 | 98 | 97.3 | 100 | 97.8 | 10.9 | 1.4 | Fc, 4 oxidation, -K/-K |
| 3 | 0.5: 99.5 | 51.7 | 46.5 | 93.3 | 84.1 | 9.8 | 2.0 | Fc, 0 oxidation,  -K/unclipped |
|  |  |  |  |  |  |  |  | Fc, 1 oxidation, -K/-K |
|  |  |  |  |  |  |  |  | Fc, 2 oxidation, -K/-K |
|  |  |  |  |  |  |  |  | Fc, 2 oxidation,  -K/unclipped |
|  |  |  |  |  |  |  |  | Fc, 4 oxidation, -K/-K |
| 4 | 0.2: 99.8 | 8.4 | 10.3 | 32.5 | 99.5 | 2.1 | 1.4 | Fc, 0 oxidation, -K/-K |
