## Supplementary material for "Characterization and Validation of a Middle-Down Hydrophobic Interaction Chromatography Method to Monitor Methionine Oxidation in IgG1": Table S2

**Supplemental Table S2:** Summary of % PTMs, bottom-up LC-MS/MS and SEC-MS analysis of purified F(ab’)2 -containing fractions. Bold indicates predominant species.

| Fraction | F(ab’)2:Fc (%) | N-terminal Pyroglutamic acid(%) | F(ab’)2 Met oxidation(%) | F(ab’)2 Trp oxidation^(^%) | F(ab’)2 His Oxidation(%) | SEC-MS |
| --- | --- | --- | --- | --- | --- | --- |
| 5 | 78.1: 21.9 | 1 | 2.6 | 18.9 | 1.5 | (F(ab’)2)( F(ab’)2) |
|  |  |  |  |  |  | (F(ab’)2)(Fc) |
|  |  |  |  |  |  | **F(ab’)2 + 11 Da** |
|  |  |  |  |  |  | F(ab’)2 with N-terminal truncation |
|  |  |  |  |  |  | Fc |
| 6 | 98.8: 1.2 | 0.3 | 1.2 | 8.1 | 0.3 | **F(ab’)2** |
| 7 | 87.5: 12.5 | 11.3 | 3.5 | 31.7 | 0.6 | **F(ab’)2 (with shoulder indicating a lower mass)** |
|  |  |  |  |  |  | (F(ab’)2)(Fc) |
| 8 | 90.3: 9.7 | 2.2 | 3.3 | 21.9 | 0.6 | **F(ab’)2 (multiple compact conformations)** |
