## Supplementary material for "Characterization and Validation of a Middle-Down Hydrophobic Interaction Chromatography Method to Monitor Methionine Oxidation in IgG1": Fig. S1

**Supplemental Figure S1:** Fraction 3 analyzed by SEC-MS. Quadruple oxidation variant with fully clipped Lys. Double oxidation variant (Met252/Met252, Met428/Met428, or Met252/Met428) with fully clipped Lys. Singly oxidized variant (Met252 or M428) with both C-terminal Lys clipped. Non-oxidized variant with one C-terminal Lys clipped.

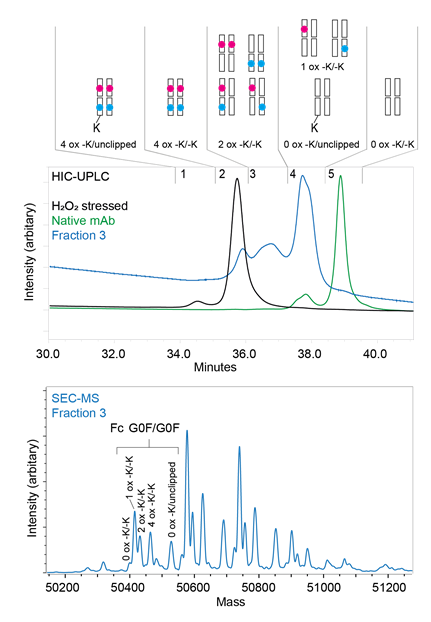
