## Supplementary material for "Characterization and Validation of a Middle-Down Hydrophobic Interaction Chromatography Method to Monitor Methionine Oxidation in IgG1": Fig. S2

**Supplemental Figure S2:** Accuracy profile of the HIC method. Relative β-expectation tolerance limits (%) per level: [-11.1%, 8.7%] for 3.771% of Fc Ox, [-7.0%, 5.6%] for 7.541% of Fc Ox, [-3.4% , 3.4%] for 15.082% of Fc  Ox , and [-2.5% , 5.1%] for 37.705% of Fc  Ox.

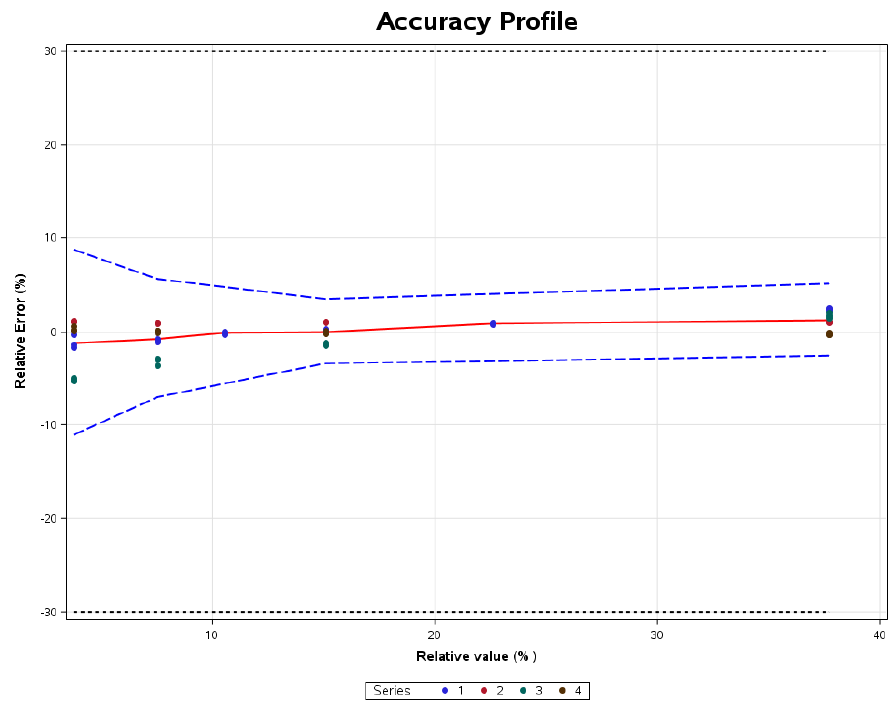
