## Supplementary material for "Characterization and Validation of a Middle-Down Hydrophobic Interaction Chromatography Method to Monitor Methionine Oxidation in IgG1": Fig. S3

**Supplemental Figure S3:** Middle-down HIC analysis of H_2_O_2_-stressed mAb after IdeS digestion. (A) MAbPac HIC. (B) Protein-Pak HIC. (C) AdvanceBio HIC.

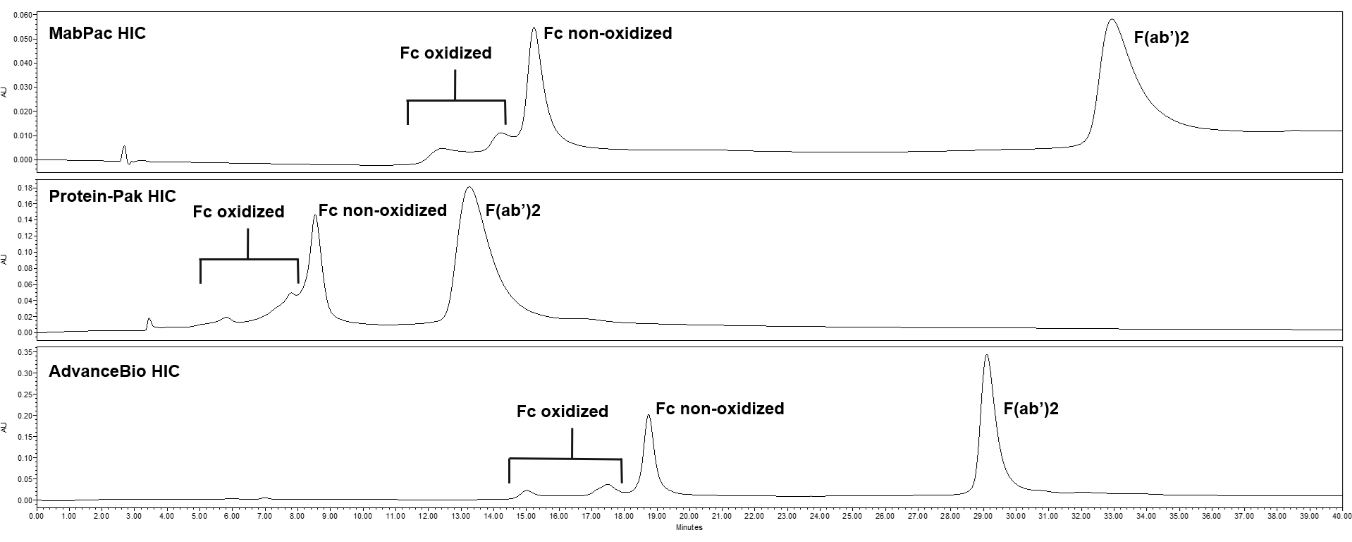
